# Conserving amphibians in the face of land development: integrating field experiments as a planning tool

**DOI:** 10.1101/183343

**Authors:** Alexander J. Felson

## Abstract

Regulations designed to guide development practices inadequately reflect ecological understanding and fall short of preserving viable habitats. Environmental consultants use rapid assessments and monitoring on individual ponds to rank pond habitat quality, relying on coarse proxies, including vegetative indicators, soil characteristics, hydroperiod, and breeding evidence in obligate species. Planners incorporate these rankings to inform the layout of neighborhoods, roadways, infrastructure and housing. However, important drivers of amphibian survival and fecundity—including metapopulation dynamics, habitat connectivity, watershed health, terrestrial density dependence, and environmental gradients—are often poorly measured and regulated. Given that development proceeds regardless, what options exists improve land development practices?

Integrating experimentation into the planning process can inform land development and improve amphibian conservation. Working as part of the design team we employed an adaptive approach called *designed experiment* to inform development practices. We manipulated *Ambystoma opacum* larvae within enclosures to test the effects of inter-pond conditions (versus intra-pond density dependence) on the survival and fecundity of conspecifics, *Rana sylvatica and Ambystoma maculatum*. While the A. maculatum populations were decimated with only 1.5 % survival. For *A. opacum* and *R. sylvatica* results indicate habitat variation between ponds accounted for 63.7% and 50.3% of the variance in survival rates of larvae, respectively, and are not predicted by the presence and abundance of egg masses, while density effects accounted for 3.5% and 2.8% of the variation in survival. The results suggest that ponds ranked as high value based on egg mass counts may actually function as habitat sinks. This study illustrates the potential value of assessment approaches that emphasize habitat quality across pond clusters to guide mitigation, conservation, regulations, and to establish sites and funding for ecological research.

## Introduction

Amphibians are declining and are especially vulnerable to land use impacts (Collins and Crump 2009) and urbanization (Baldwin and deMaynadier 2009; Scheffers and Paszkowski 2012). Their complex life cycle—including an aquatic larval stage, metamorphosis, and a terrestrial-aquatic adult stage—expose amphibians to development, which often damages their upland or aquatic habitats and connectivity (Semlitsch 2002; Gamble et al. 2006) and exacerbates the naturally stochastic population fluctuations that arise from seasonal and inter-annual variability and climate change (Collins and Crump 2009). Land use change is a leading cause of amphibian declines. Regulations, however, inadequately reflect ecological understanding and fall short of preserving viable habitats (Snodgrass et al. 2000). Many factors impacting amphibians are unregulated including fragmentation of pond habitats, seasonal disruption of watersheds (Windmiller and Calhoun 2008), upland habitats (Becker et al. 2010), pond networks, and metapopulations (Petranka 2007). This is partly due to the challenges of translating scientific understanding into policy as well as the inflexibility of the legal framework to accommodate shifts in ecological perspectives (McGreavy et al. 2012). Ecologists have proposed additional guidelines (e.g. Calhoun et al. 2005) however, land use law and the legal realities surrounding property ownership tend to favor property owners over wildlife and there is no incentive to go beyond the regulations (Gamble et al. 2006).

The complex biphasic life cycle patterns of amphibians, combined with the inherent difficulty of monitoring fossorial species, challenge efforts to establish laws and management practices that respond to amphibian needs (Blaustein and Kiesecker 2002). Indeed, vernal pool regulations have proven ineffective in mitigating development impacts (Mahaney and Klemens 2008). In all cases amphibian assessment informing development tends to focus on the aquatic stage density dependence based on coarse and rapid assessments of species presence and densities within ponds (Cowardin 1979). Past experiments mostly focused on individual ponds (Petranka and Sih 1987; Pechmann et al. 1989), and likely further reinforced assumptions that the presence of diverse populations within individual ponds indicates which merit conservation. However, the results of long-term research (Werner et al. 2007; Berven 2009; Gamble et al. 2009) and recent studies challenge these assumptions. Researchers now acknowledge the importance of additional factors including: terrestrial conditions (Shulse et al. 2010; Van Buskirk 2010) habitat fragmentation (Becker et al. 2010; Petranka 2007), and terrestrial stage density dependency linked to juvenile survival (Berven 2009). Additionally, in view of the stochastic nature of habitat characteristics (i.e., the feedback between rainfall, hydroperiod and population dynamics), even poorly performing ponds in a given year may still play an important role in longer-term species survival (Petranka 2007; Pechmann et al. 1989). Improving the evaluation of development activities will require better rapid assessments, improved metrics, better integration of science with policy (McGreavy et al. 2012), and multi-year assessments across pond clusters to capture stochastic patterns (Semlitsch 2002).

Recognizing regulations change slowly, we sought an alternative model, called *designed experiments* (Felson and Pickett 2005; Felson et al. 2013), to position ecological research as an integral component of the planning. Researchers worked as part of the design team with the support of the developer to assess pond migration patterns to guide site design. The *designed experiment* provided the developer the means to negotiate with the planning board and proactively avoid future postponements (Felson 2007). The designed experiments approach additionally offered: (1) a platform to situate hypothesis-driven research on a development parcel, a land use historically inaccessible to ecologists; (2) a method for generating robust scientific data to expose ecological processes; (3) a cross-disciplinary framework for experiential learning and an iterative process connecting ecologists with developers, consultants, planners and officials to refine the design.

## 1 Materials and Methods

### 1.1 Study site

Tuxedo Reserve is a 500 ha parcel along the Ramapo River 68 km northwest of New York City and adjoins two protected areas, Harriman and Sterling Forest State Parks. The mostly forested parcel is 6% forested wetland and 1% vernal pools. In the spring 2008 the pool size ranged from 0.067 to 0.89 ha, with mean depth ranging from 0.16 to 0.38 m. Fed by overland and groundwater flow, the ponds dried out between June 26 and July 21. Habitats are predominantly red maple hardwood swamps but include two shrub bogs.

### 1.2 Endemic amphibian populations

Vernal pools in the Reserve support multiple amphibian species. *Ambystoma opacum* (marbled salamanders) is a dominant, fall-breeding species. *Rana sylvatica* (wood frogs) and *Ambystoma maculatum* (spotted salamanders) are potential prey for *A. opacum* and both are common across the east coast and on the site. All three indicator species require vernal pools for reproduction. We selected these amphibians because together they provide a guild for assessing species interactions within enclosures as well as assessing the impacts of pond conditions on survival and fecundity.

### 1.3 Environmental assessments conducted on the site

The town planning board had granted preliminary subdivision approval for the 1375-unit development project; however, the board requested the developer reduce encroachment on vernal pools (Tuxedo Reserve Owner LLC September 2009). To address those concerns multiple Environmental Impact Statements were prepared (1999, 2003, and 2008) For the Environmental Impact Statement, EcolSciences Inc. delineated vernal based on four criteria: (1) occurrence in a depression with no permanent outlet; (2) breeding evidence from obligate fauna; (3) water retention during a normal rainfall year for two or more contiguous months between March and September; and (4) void of fish or dried out during a normal rainfall year (Fennessy et al. 2009).

Field studies conducted in the spring of 2006, 2007 and 2008 ranked the ecological value of isolated wetlands based on presence of Species of Special Concern, productivity, and diversity. Call recognition was used for wood frogs; egg masses were used to identify spotted salamanders, marbled salamanders, and wood frogs. Several Species of Special Concern were found: *A. opacum, Ambystoma jeffersonianum* (Jefferson salamander), *Carphophis amoenus* (worm snake), and *Clemmys guttata* (spotted turtle) (Tuxedo Reserve Owner LLC September 2009).

Following these initial efforts our team was invited to set up both applied and basic science experiments across the site to guide the development layout. Working with the developer, we first established migration studies to address concerns over road and housing proximity to wetlands (Fig. 1). We expanded this study to 5 ponds in the second season. In the third season we tested the efficacy of the rapid assessment methods used previously with a comparative larval density study across ponds. Results from the third season are the focus of this paper.

**Figure 1.**
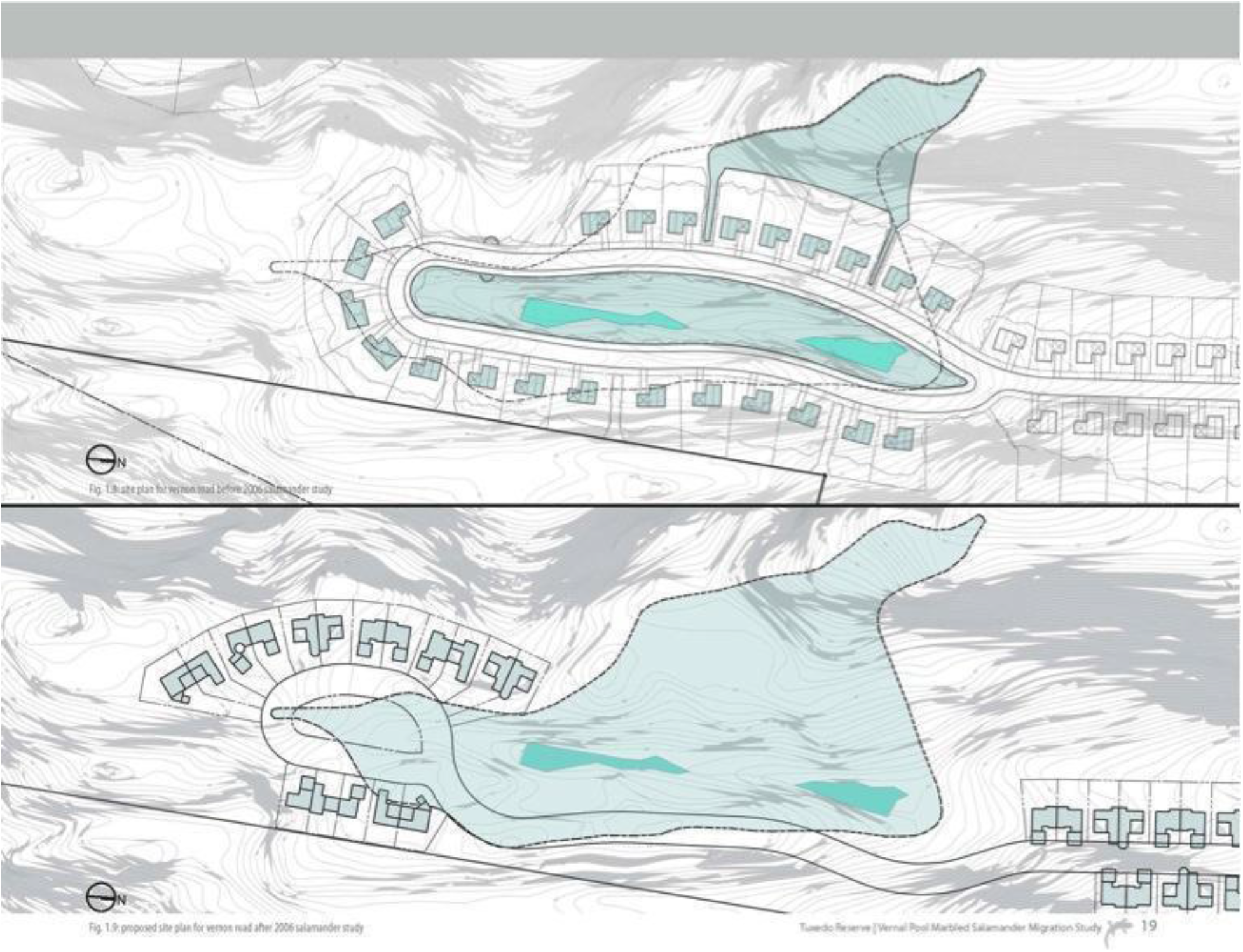
A drift fence study revealed directional migration paths. The study was used to guide decisions regarding road and housing layout.

### 1.4 Experimental design

A larval density study was set up using mesocosms across 8 ponds including several ponds previously identified by the consultants as “high value”, and one pond off-site to establish a baseline and control. This study intended to assess the effectiveness of the consultants’ pond ranking practices and further inform decisions about siting development.

We established the hypothesis that survival and fitness as measured by growth rate and fecundity will exhibit an inverse relationship to *A. opacum* densities. To compare the effects of habitat quality *across* ponds to the effects of predator density *within* ponds, we manipulated *A. opacum* counts in enclosures within and across ponds and monitored survival, size, and growth rate (following Wilbur 1997). We first manipulated the larval counts of *A. opacum* within enclosures to simulate inter-annual variation; we then assessed density effects on the survival of conspecifics and *A. maculatum* and *R. sylvatica*, across ponds. While the project focused on species performance, we did examine several additional physical and biological characteristics including: hydroperiod, depth, adult sex ratio, mean larval period, and diversity (Fig. 2). Our findings indicate pond conditions, and their associated habitats, are primary factors determining which ponds are of highest conservation value.

**Figure 2.**
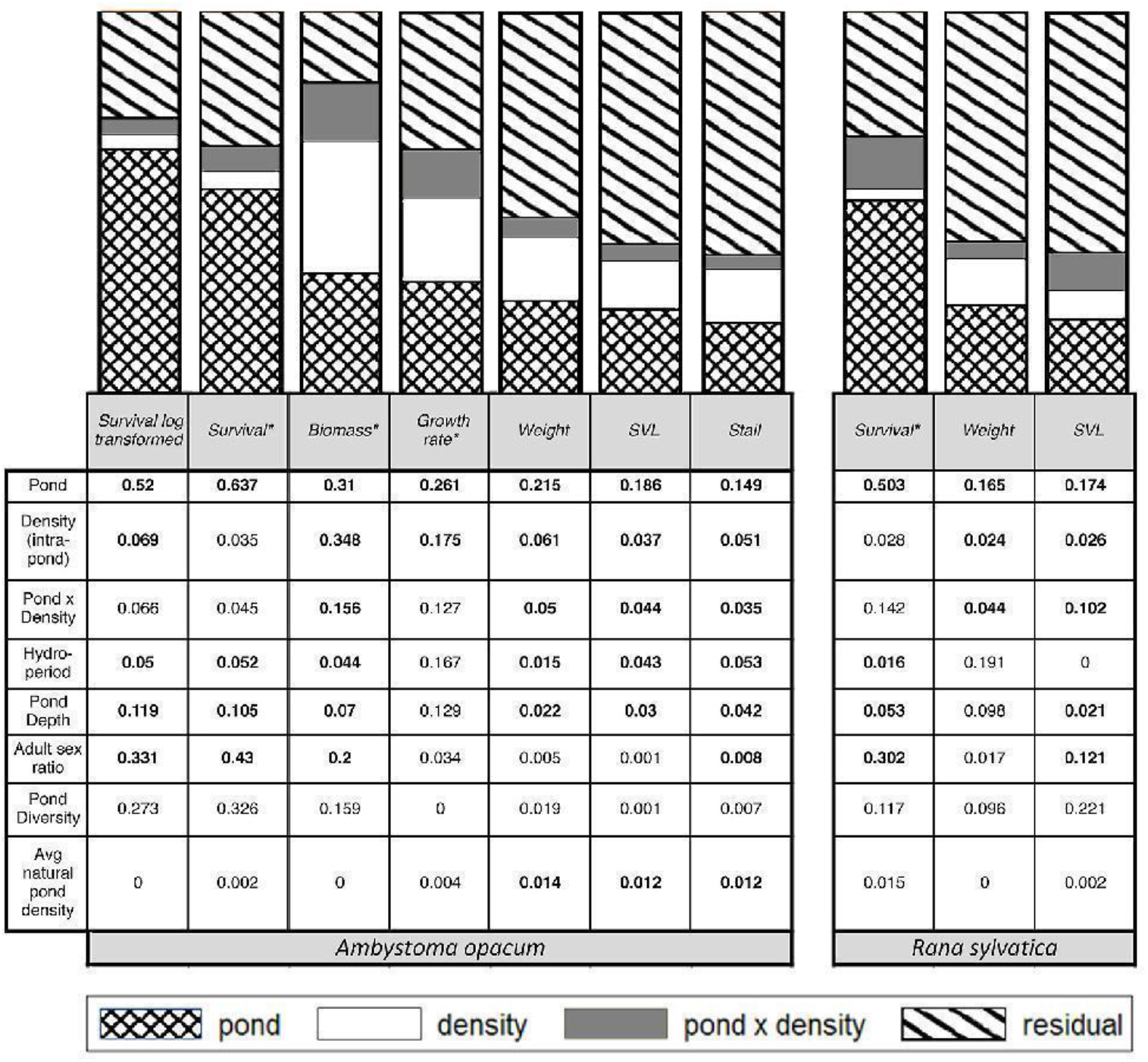
Summary of R-squared results. All independent (y-axis) and dependent variables (x-axis) with significant p-values are bolded. An * indicates variables that were measured at the enclosure scale. The bar chart above indicates variance explained by pond, density and pond x density for the dependent variables.

Within each enclosure, we varied the initial density of A. *opacum* between absent (0), low (8), medium (16), high (24) density while keeping prey species densities constant with 24 *A. maculatum* and 24 *R. sylvatica*. In determining species counts, we considered the relative abundances of larvae and egg masses in ponds as well as past experiments (Morin, 1995; Wilbur 1997). The density range of *A. opacum* was intended to mimic population fluxes across annual breeding events (Scott 1990). Each pond included 12 enclosures with three replicates (low, medium, high, and absent) *A. opacum* densities. The total number of enclosures was 96.

Enclosures had external dimensions of 60 x 60 x 240 cm, for a volume of 0.864 m^3^ (Morin 1986). Given variations in depth, water volume in the enclosures ranged from 0.23 m^3^ to 0.69 m^3^. Pens were constructed using 12.7 cm PVC pipe and fiberglass window screening (7 meshes/cm). Screens were wrapped on all sides of the enclosures and secured with greenhouse PVC clamps which allowed indigenous prey (e.g. zooplankton and insects), predator larvae (e.g. leeches, beetles, bugs, dragonflies, damselflies, caddisflies) and parasites (e.g. bacteria) to circulate freely through the cages while excluding large invertebrate predators (e.g. birds, snakes or raccoons) (Gibbs et al. 2007). Leaf matter collected from the adjacent forest provided 8 cm of substrate for refugia and moisture retention. Placing pens perpendicular to the pond edges provided deeper water to dry land transitions for refuge during metamorphosis.

To assess productivity and fitness of *A. opacum* we measured survival, average change in weight, and average change in snout-to-ventral length (SVL) (Fig. 2). For other species, we measured survival and mean biomass.

To assess habitat quality, we examined the effects of hydroperiod, depth, diversity, pond size, dissolved oxygen, salinity, and conductivity. We also assessed adult sex ratio, mean larval period, and average natural density of *A. opacum*. To evaluate larvae composition and determine mean densities, we randomly laid out four parallel transects in each sample pond between March 29 and May 22, 2008. A bucket with the base removed and a diameter of 0.54 m was inserted into two locations along each transect. We used small nets to skim the surface of each enclosure for species above the leaf matter and a dip net to disturb and sample species beneath the leaf matter, sampling with five sweeps of each net.

To avoid logistical challenges and spread of disease, we maintained larvae and egg masses within their natal ponds rather than redistributing them. We collected *A. opacum* larvae in April and combined them to randomize their genetic variation and hatch dates within individual ponds. In addition, *A. maculatum*, and

*R. sylvatica* egg masses were counted within each pond and we used a gridded white background to photograph and count the number of eggs per mass. Enclosures were stocked using a subsample of the egg masses. There were two exceptions: *R. sylvatica* from pond 5 were used in pond 2, and *A. maculatum* from pond 2 were used in pond 1, because viable populations were not available in those ponds.

Monitoring throughout the summer ensured enclosures were free from outside predation. Enclosures were shifted down-slope into the remaining wet areas as water levels decreased. As each pond dried out, we took a complete census of survivors, including weight and length.

### 1.5 Effects of pond habitat quality and initial predator density on performance

A two-factor MANCOVA controlling for initial weight was used to determine whether differences across ponds, manipulated predatory larval densities, or the interaction between individual ponds and predator densities impacted species performance. The organism-based performance variables included: larval survival (percentage of the initial number stocked collected after metamorphosis), Arcsine survival, snout-to-vent and snout-to-tail length at metamorphosis, mean wet mass at metamorphosis, biomass, and a linear approximation of mean growth rate per day (the difference between mean metamorph body size and initial mean body size for *A. opacum* /mean larval period). The three covariates in the MANCOVA were: mean initial weight, snout-to-vent length, and vent-to-tail length. Because mean predator size initially differed between ponds and different predator densities, we controlled the initial predator density using covariates for mean initial *A. opacum* size.

Survival rates were regressed on habitat quality and predator density (Fig. 2) using indicator variables of pond and density. *A. opacum* density was measured and tested as a categorical and a continuous variable. The effects of pond-level variables on measures of performance were analyzed using scatterplots. The output weight and size of *R. sylvatica* were evaluated although the growth rate could not be determined because the initial tadpole sizes were too small to record. Summary response variables for *A. opacum* included survival, mean larval period, mean weight, length and biomass at metamorphosis, and growth rate. Our results from testing the effects of pond-level variables on other measures of amphibian performance similarly explained little of the variation.

## 2 Results

### 2.1 Ambystoma opacum survival

Survival of *A. opacum* varied dramatically across ponds (Fig. 3). A. *opacum* survival rates ranged from over 80% in ponds 4 and 8 to less than 5% in pond 6. Several ponds perform similarly to one another. An ANOVA revealed that 63.7% of the variance was between ponds (F=17.23 and p<0.001) (Table 1). Generally, survival of *A. opacum* was inversely related to initial predator density (Fig. 4). According to the ANOVA analysis, the effect of initial predator density on survival was significant (p<0.05), but accounts for only about 3.5% of the variance in *A. opacum* survival.

**Table 1.**
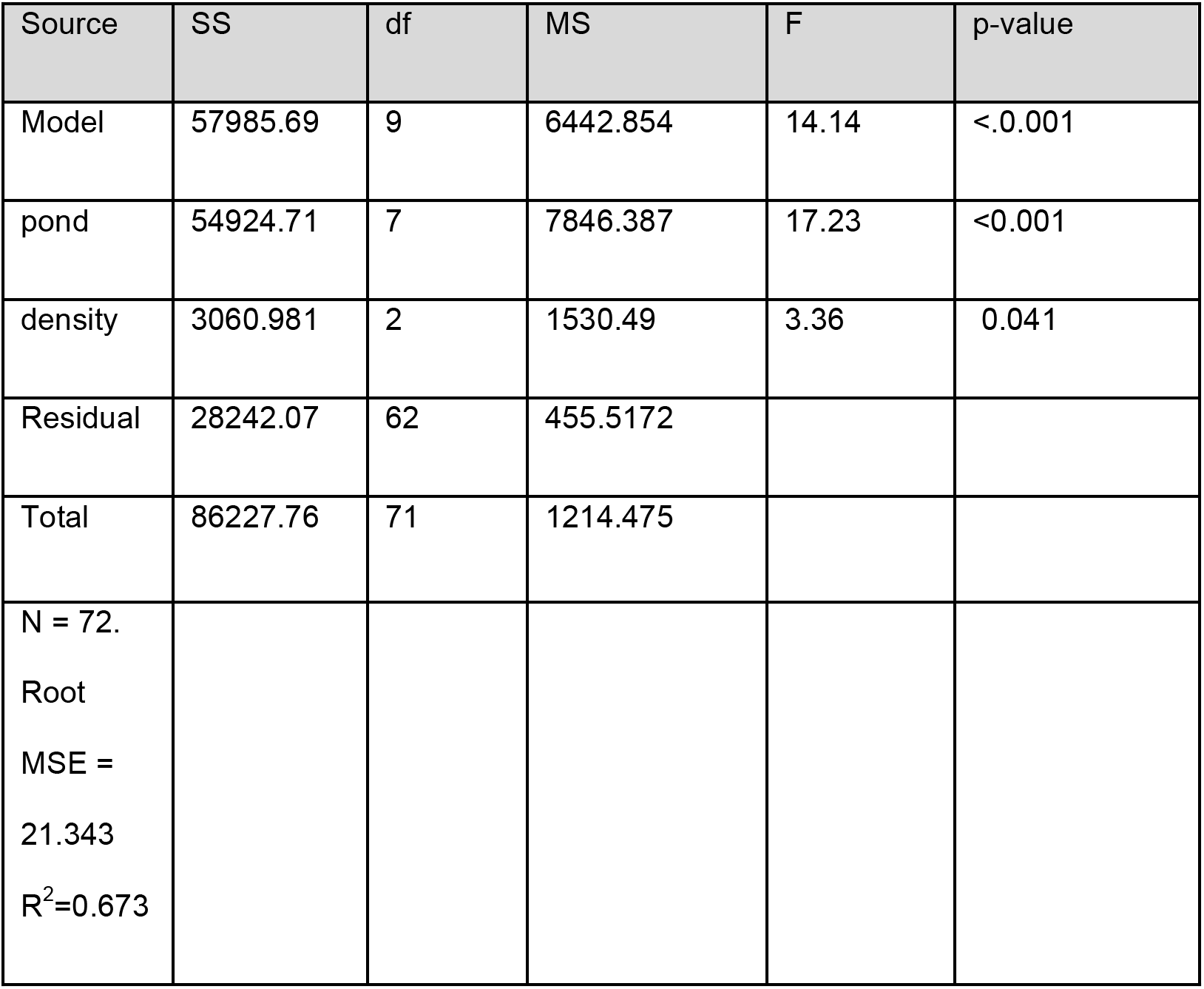
ANOVA predicting Amybstoma opacum survival rate depending on pond and density.

**Figure 3.**
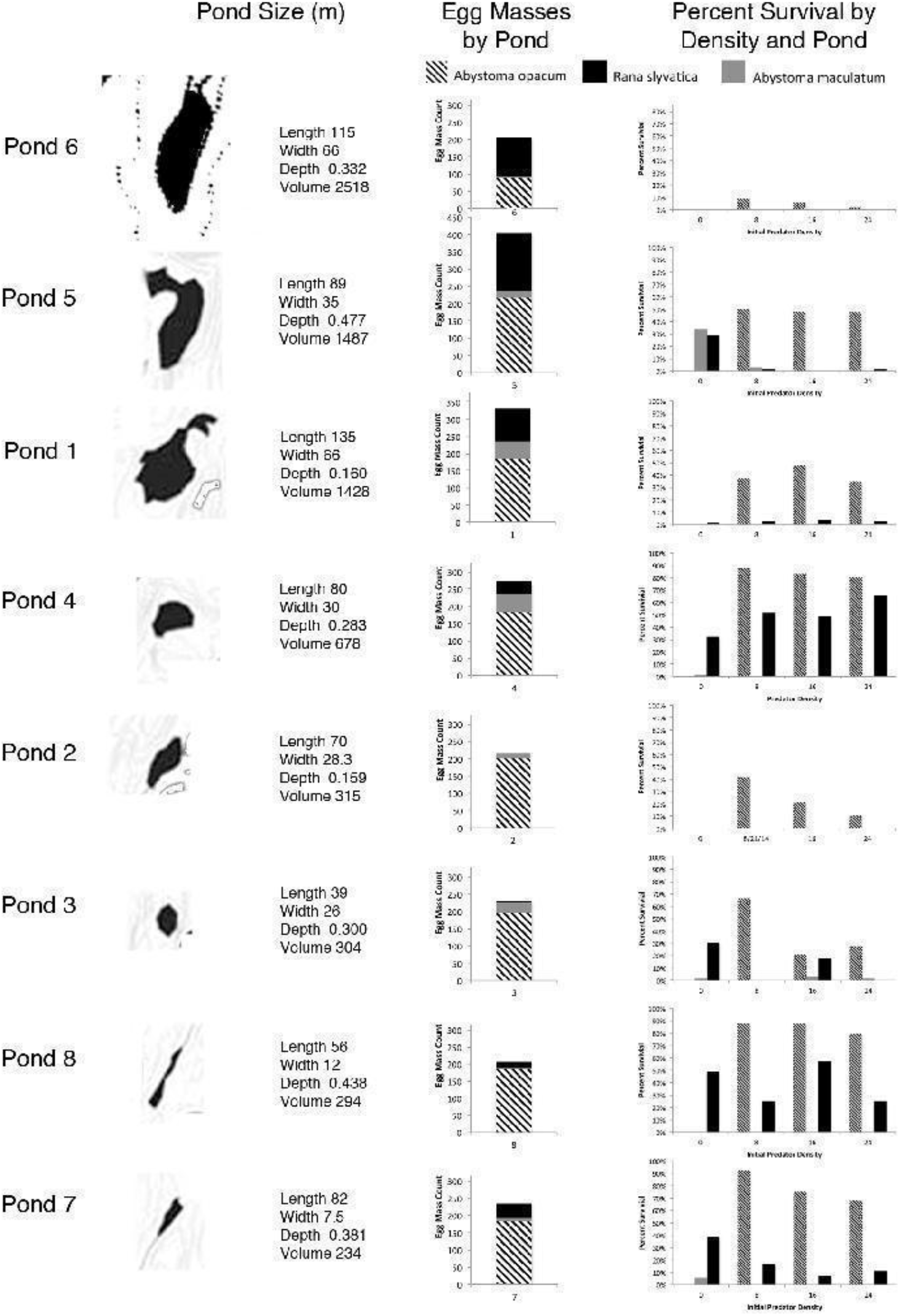
Ponds are shown ranked by volume. Egg mass counts, shown in the center column, were used as an initial predictor of pond habitat value. Egg mass counts were not a good predictor of survival shown in the right column.

**Figure 4.**
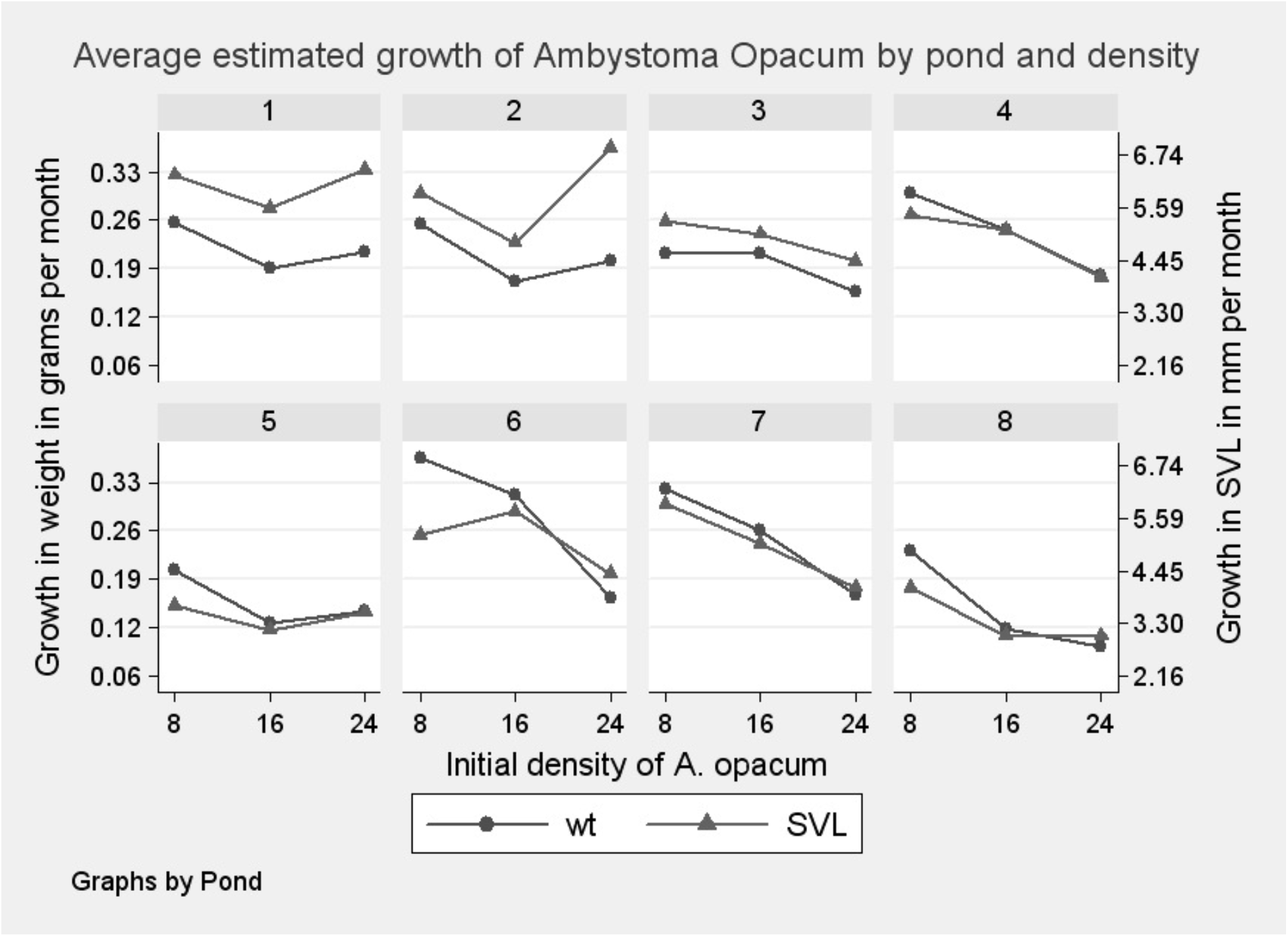
Graphs illustrating average estimated growth of *A. opacum* by pond and density.

### 2.2 Ambystoma opacum size

Because the initial size of *A. opacum* individuals differed, we used a two-factor MANCOVA analysis controlling for initial weight to analyze output size (Table 2). The mean output weights of *A. opacum* populations ranged from 0.65 grams to 1.05 grams, while the mean length ranged from 25.8 mm to 34.5 mm (Fig. 2). In both cases, pond 6 has the highest average weight and length, likely because of decreased competition for food. The variation in *A. opacum* weight across the remaining ponds was considerably lower.

**Table 2.** MANCOVA predicting weight, snout-to-vent length and tail length for *Ambystoma opacum*. Lower values of Wilks’ Λ indicate larger effects and MANOVA of *Ambystoma opacum* Growth Rate. MANOVA/MANCOVA of *Rana sylvatica* weight and length (excluded enclosures where *A. opacum* initial density equaled zero).

| MANCOVA *Ambystoma opacum* length and weight | F* | num. df | den. Df | P-value |
| --- | --- | --- | --- | --- |
| *A. opacum* Initial Mean weight | 7.6 | 1 | 37 | 0.0017 |
| Pond | 2.67 | 7 | 74 | 0.0033 |
| Density | 13.21 | 2 | 74 | <0.001 |
| Pond x Density | 1.13 | 14 | 74 | 0.3312 |
| MANOVA *Ambystoma opacum* growth rate |  |  |  |  |
| pond | 7.84 | 21 | 106.79 | <0.001 |
| density | 3.73 | 6 | 74 | 0.003 |
| pond x density | 1.4 | 42 | 110.53 | 0.084 |
| MANCOVA *Rana sylvatica* weight & length<br>controlling for initial *A. opacum* weight |  |  |  |  |
| mean initial AMOP weight | 7.07 | 2 | 222 | <.001 |
| pond | 53.54 | 21 | 106.7941 | <.001 |
| density | 7.40 | 6 | 74 | 0.003 |
| pond x density | 4.23 | 42 | 110.5251 | 0.084 |
| MANOVA *Rana sylvatica* Weight and length |  |  |  |  |
| Pond | 53.97 | 6 | 654 | <0.001 |
| Density | 5.59 | 6 | 654 | <0.001 |
| Pond x Density | 4.68 | 14 | 654 | <0.001 |

\* Wilks' lambda was used for F tests.

After adjusting for initial differences in mean weight, MANCOVA analysis revealed significant effects of pond habitat conditions, predator density, and the interaction between pond habitat and predator density on A. opacum size (Table 2). Average size consistently declined as initial predator density increased in ponds 4, 5, 7, 8, although this decrease was not significant in pond 5 (Fig. 1). Average size did not consistently decline in ponds 1, 2, 3 and 6. Thus, interaction in the MANCOVA results between pond habitat and predator density reveals that predator density impacts A. opacum size in some, but not all ponds (Table 2).

### 2.3 Impact of Initial Predator Density on A. opacum size

A regression showed *A. opacum* weight varied significantly across predator densities. Where there was initially a low density of predators mean *A. opacum* weight was 0.856 grams, whereas at medium density it was 0.749 grams and at a high density, 0.665 grams. At high density *A. opacum*, there was minimal variation in weight gain, although length had a greater range. Survival of *A. opacum* did not significantly differ for low and medium densities. These results imply initial predator densities only weakly influence performance of amphibians.

### 2.4 Ambystoma opacum growth rate

Growth rate of *A. opacum* larvae was determined using mean size and length of larvae within each enclosure. Pond effects accounted for 6.9% of the growth rate for A. opacum. Growth rates both decreased and increased with higher initial predator densities. In ponds 3, 4, 6, 7, growth rates decreased from 2.73 mm (low) to 2.58 (medium), and to 2.00 (high) as predator densities increased. In ponds 1, 2, 5, 8 average growth rate decreased from 3.02 (low) to 2.11 (medium) when predator density was increased from low to medium, but increased from 2.11 (medium) to 2.18 (high) when density was increased.

MANOVA analysis reveals significant effects of habitat conditions and density on growth and survival of *A. opacum*, but the interaction between habitat conditions and density is not significant. In addition, the Wilks’ Λ value for the effect of habitat quality on growth is 0.069, whereas the Wilks’ Λ value for the effect of predator density on growth is 0.589. This indicates that the effect of habitat quality is 8.5 times greater than the effect of initial predator density on *A. opacum* growth (Table 2).

### 2.5 Rana sylvatica survival

Survival of *R. sylvatica* varied substantially across ponds ranging from no survivors to 18.4% in pond 7. Because *R. sylvatica* individuals died out in more than half of the enclosures, we used a logistic regression to predict survivorship. A linear regression was used to predict whether *A. opacum* density would significantly affect the number of *R. sylvatica* survivors within enclosures in which there was at least one survivor. Based on 95% confidence intervals, initial *A. opacum* density did not affect survival of *R. sylvatica* in any of these models (Fig. 2). Although survival rates of *R. sylvatica* were 23% in enclosures where *A. opacum* were absent and 14% in other enclosures, the difference is not significant. Moreover, the survival rate of *R. sylvatica* was 12% when grouped with low *A. opacum* and 13% when grouped with high *A. opacum*. Pond effects accounted 50.3% of the variance in survival rate of *R. sylvatica*.

### 2.6 Rana sylvatica size

Because of their small size, we did not measure initial size of *R. sylvatica* individuals and nor calculate growth rate but did determine output size. Measurements were limited to ponds 3, 4, 7, 8 because there were very few *R. sylvatica* survivors in ponds 1, 2, 5, and 6.

As the density of *A. opacum* increased, output weight and size of *R. sylvatica* increased significantly in ponds 4 and 8 from 0.374 g to 0.533 g (p<0.001) where initial predator densities were absent and high, respectively. Snout-to-vent measurements were 15.53 and 17.07 mm (p<0.001) where predator densities were absent and high. Weight and size of *R. sylvatica* did not significantly increase with density in ponds 3 and 7 (Fig. 1). Habitat quality, predator density, and the interaction between pond and predator density all significantly affected weight and length of *R. sylvatica* (p<0.05).

Random differences in initial *A. opacum* weight are positively associated with the final weight of *R. sylvatica*. A Tukey test revealed *R. sylvatica* weight is significantly different between ponds 1 and 8, although *R. sylvatica* weight and length does not vary between ponds 4 and 7. For *R. sylvatica* length, the Tukey test indicated all differences are significant except between ponds 4 and 8. The particularly low weight in pond 8 distinguishes it from ponds 3, 4, and 7, which are more tightly clustered. In sum, the lower values of Wilks’ Λ indicate pond habitat conditions affect *R. sylvatica* weight and length more than initial predator density or initial sampled population density.

### 2.7 Additional factors across ponds

To account for habitat quality, we documented the effects of hydroperiod, adult sex ratio, mean larval period, depth, size and diversity of ponds, and average natural density of *A. opacum*. We also returned to measure dissolved oxygen, salinity, and conductivity in spring 2009. The effects of these pond-level variables were evaluated against the summary dependent variables including survival, weight, length, and growth rate (Fig. 2).

Hydroperiod causes stochastic population fluctuations in individual pond communities over time (Snodgrass et al. 2000). Because our study spanned one season, the same weather patterns affected all ponds. We compared ranges of hydroperiod across individual ponds in a given season. Ranges in hydroperiod are likely attributable to differences in the physical and biotic conditions of ponds and their watersheds. We found the range of the hydroperiod was significant (explaining about 5.8% of the variability) for all of the *A. opacum* dependent variables except growth rate, but only for survival of *R. sylvatica* (Fig. 2). However the inter-pond variation and within pond enclosure survival, size, and growth rate did not appear to be driven by hydroperiod. Differences across ponds did not vary enough to alter hydroperiod in a given year, nor affect performance.

Pond depth proved significant for survival but not for size of *R. sylvatica* and for all of the *A. opacum* dependent variables except growth rate. Adult sex ratio proved significant for survival for growth rate, weight and length (SVL) (Fig. 2). The relationship between *A. opacum* survival and adult sex ratio appears to be positive. Low R-squared values indicate that neither factor explains most of the variation linked to pond effects. Pond depth explained about 7.6% of the pond variation. However, the pond performance does not seem to be driven by these variables and the low n value (n=8) makes it challenging to disentangle the influence of independent variables. Thus, most of the variation in ponds remains unexplained.

Other qualitative factors including the presence of predators, specifically leeches and Dragonfly nymphs, were observed in ponds 1, 2, and 6. Their presence may explain the low survival rates in these ponds.

## 3 Discussion

In line with previous work (Werner et al. 2007; Veysey et al. 2011) our research on migration and larval densities across multiple ponds, provides additional evidence of the importance of landscape scale habitat assessments. We assessed varying population densities within ponds with the goal of improving habitat assessments guiding suburban planning. We determined the extent to which population density and differences among habitats played a role in survival and growth. Similar to recent studies focused on the impact of canopy cover gradients on organism distributions across ponds (Van Buskirk 2010), and building on previous studies concentrated on intra-pond performance (Scott 1990; Lawler and Morin 1993), our study focused on comparing the impact of varying population density within ponds to the impact of habitats on performance. Our findings reveal considerable variation amongst breeding populations across ponds in terms of survival, size, and growth rate. Pond habitat conditions rather than population densities appear to be the dominant factor impacting survival (Fig. 2).

However, the current habitat assessment practices emphasize species presence over extensive habitat assessment. Habitat quality is assessed rapidly using a wetland classification systems such as the Cowardin Classification system, which defines wetlands based on plants, soils, and flooding frequency (Cowardin et al. 1979). Other factors, such as canopy cover or wetland gradients are not considered, despite evidence of the importance of these physical traits (Werner et al. 2007; Skelly et al 2013). Wide buffers, watersheds and pond interconnectivity are generally inadequately integrated into conservation regulations given the importance of upland habitat quality, metapopulation dynamics and water quality on performance. To improve conservation of ponds with the highest productivity and survival requires going beyond existing laws (Calhoun et al. 2005) and eventually revising them (Semlitsch 2002).

### 3.1 Initial predator density effects on performance

Our in-situ experimental design was intended to loosely mimic natural larval densities and the life cycle sequence as amphibians migrate. We based experimental larval densities on sampled densities and on previous reports of the minimum numbers needed to elicit meaningful results (Morin 1995). Our finding that predator density is not a major contributor to population performance contrasts with previous studies (Skelly and Kiesecker 2001; Altwegg 2003). Our comparison was limited to one season across multiple ponds whereas other studies focused on individual ponds over a longer term. Field and laboratory studies have shown increases in aquatic larval density result in lower survival rates, slower growth, longer larval periods, and reduced body size of the species at metamorphosis (Petranka and Sih 1987; Taylor and Scott 1997). Previous work indicated aquatic larval densities in individual ponds account for 37-89% of the growth of *Amybstoma* spp. and for less than 50% survival of *Rana sylvatica* (Semlitsch 1987; Wilbur 1987). Data on the density effects of the upland terrestrial life stages are rare, although a recent long-term study indicated a large terrestrial density effect of *R. sylvatica* on juvenile survival and population size (Berven 2009). Other researchers have emphasized the need for more research on movement, recruitment and population persistence in order to refine our knowledge of amphibian lifcycles (Scheffers and Paszkowski 2012).

In our study, initial predator density significantly affected several measures of performance. However, density only accounted for 3.7-18% of variation in *A. opacum* survivorship and 2.4-2.6% of variation in *R. sylvatica* survivorship across ponds (Fig. 2). Contrary to expectation, we found no inverse relationship between *A. opacum* density and the fitness of conspecifics, *R. sylvatica* and *A. maculatum*. Manipulated *A. opacum* densities did not significantly impact *R. sylvatica* survival (P=0.177), and survival rates actually increased with an increase of *A. opacum* density from 8 to 24 individuals. Competition and predation between *A. opacum* and *R. sylvatica* did not seem to be an important factor in determining survival of *R. sylvatica*, as the weight of *R. sylvatica* significantly increased with higher density of *A. opacum* in ponds 4 and 8. It is possible that the higher density mesocosms amplified the overall population that could host leeches, odonates and other predators or parasites leading to greater pressures on *R. Sylvatica* in higher density mesocosms.

Predator density effects may have outweighed pond effects for *A. maculatum*, which were decimated in all instances with only 1.5% survival in ponds 3, 4, 5, and 7. *A. maculatum* survival was slightly better in enclosures with no *A. opacum* (5.2%). These results have several possible explanations. 1) Stressful environmental conditions may create more pronounced negative effects of density on larvae (Werner et al. 2007); 2) Ecological effects might be overestimated as a result of unrealistically high densities in enclosures (Scott 1990); 3) *A. opacum* may have acted as either a predator or a competitor of *A. maculatum*; 4) *R. sylvatica* may have competed with *A. maculatum*. Further studies including the manipulation of prey species densities are needed to determine why *A. maculatum* failed to thrive. At the same time the small enclosures could have protected the larvae and artificially improved larval survivorship over natural survival. This may have lead to increased competition for *A. maculatum* (Scott 1990).

### 3.2 Ponds and metapopulations

The metapopulation dynamics of amphibians are often ignored by development practices that only consider individual pond preservation. On the Tuxedo site, we identified a pattern of productive and unproductive ponds, consistent with metapopulation models. Larvae did not survive in some ponds, while survival rates were high in others. A cluster of populations is not synchronized or predictable. It is probable juvenile dispersal from more productive ponds may support a cluster of less productive ones. Most breeding populations are spatially discrete; adults return to the same ponds to breed and low rates of juvenile dispersal occurs across ponds (Smith and Green 2005). Since juvenile population at breeding ponds can vastly exceed replacement rates, juvenile and adult densities commonly exceed the resource and habitat limitations of the terrestrial habitat surrounding ponds (Berven 2009). In such a situation, terrestrial density dependence may limit population size.

Our decision not to randomize the sample populations across ponds increased the genetic variability between ponds and resulted in differences in the initial size of *A. opacum*, which was controlled for in the analysis. Maintaining discrete populations revealed patterns of pond-dependent survival and growth, consistent with metapopulation dynamics. Development practices that only consider preservation of individual ponds rather than entire clusters are missing important landscape scale patterns. As researchers have recognized, our ecological understanding can surpass incorporation of this information into regulations and planning (Shulse et al. 2010; Scheffers and Paszkowski 2011).

### 3.3 Pond Size

Although our results reveal substantial effects of pond characteristics on performance, the measured variables do not completely explain differences between ponds. A number of causal factors may be confounded, including biotic and abiotic components as well as the genetic and biological differences between populations. Pond size might explain differences between ponds although for our study this was not a main contributor predicting survival or growth rates. With the exception of pond 2, the larger ponds 1, 5, 6 did not perform as well as the smaller ponds 3, 4, 7, 8 with respect to *R. sylvatica* and *A. opacum* survival, growth rate and size. This contrasts with a common assumption that larger wetlands are inherently more valuable. New York regulations, for instance, ignore ponds under 5 ha (Mahaney and Klemens 2008). However, smaller, less diverse ponds have been shown in some cases to have higher productivity than larger, more diverse ponds (Snodgrass et al. 2000). Our data is not inconsistent with the idea that pond size explains differences in survival and growth; however, with only 8 ponds it is not possible to definitively conclude its importance.

### 3.4 How effective is habitat quality monitoring?

Consultants predominantly evaluate ponds by rapidly assessing flora, fauna, substrate, and hydrology (Cowardin et al. 1979). The presence and quantity of egg masses and larval species as well as call recognition are used to rank habitats (van Horne 1983). Our research indicates that the presence of egg masses does not distinguish productive ponds from those that receive egg masses but function as sinks and undergo periodic extinctions. In another study the size and quantity of eggs in a pond reflected the survival and density of terrestrial breeding populations more than pond quality, when at lower juvenile populations, females produce smaller eggs in greater number than at higher juvenile populations (Berven 2009). Thus, egg masses likely reflect a combination of suitability of pond habitat, population dynamics, and the impact of terrestrial conditions on pond life.

Counting egg mass makes sense as a rapid and generalized approach that provides conservative results; however, in this case ponds with high population densities illustrated both high and low survival and fecundity. Egg mass data could be misinterpreted to suggest that we can preserve habitat on a pond-by-pond basis. The method does not predict the likelihood of hatched larvae to thrive. Reproductive recruitment surveys such as this study can improve accuracy. Based on our findings, parameters used by consultants to predict pond value should expand to further evaluate pond and surrounding habitat quality.

Regulations guiding amphibians conservation do not adequately reflect our understanding of the species’ life cycles. Fundamental factors including preserving upland habitats or migration routes, watersheds and the interconnectivity of pond clusters (Petranka 2007), all fall outside the regulations. Therefore, even regulated vernal pools are unlikely to facilitate the maintenance of metapopulations. In the U.K. regulations explicitly protect only a limited number of species and habitat destruction remains a primary contributor to amphibian decline (BeeBee 2014). In the U.S. the depth of ecological analysis required under NEPA can vary substantially from one environmental impact assessment to the next (Gibbs 2005).

## 4 Opportunities for designed experiments in planning projects

Recognizing gaps exist between ecological science and regulations, we established *designed experiments* that exceeded regulations and applied ecological research to the design process. Consultants working for developers often have no choice but to advocate for preserving certain ponds and sacrificing others. Research-based approaches can therefore help consultants make more informed decisions about how to couple large-scale development with efficient preservation (Felson et al. 2013).

Based on these studies we recommend: (1) framing research as a proactive tool that can support basic science and produce more specific information about where to build. (2) building on the flexibility of the legal system at the local level to influence comprehensive plans and zoning regulations and (3) creating a direct link between ecologists and practitioners to ensure that the design choices reflect the best scientific understanding. In this way, the ecologist can play a key role in defining habitat preservation and can guide projects based on site-specific research. This approaches advances the relationship between ecology and development where ecological information typically only influences design indirectly through regulations.

## 5 Conclusions

We propose a new strategy for integrating ecological research into development projects by creating active collaboration between researchers, developers, and designers. Our approach acknowledges that amphibian populations are not likely to succeed under current regulations. We therefore propose proactively inserting research into the design of a development. By integrating research into the project the design team was able to go beyond the current regulations and improve the potential for survival. Based on our findings, conservation efforts should focus more on pond and upland habitat quality with assessments of diversity, abundance and composition of populations. Conservation could be more effective if growth and survivability of target species is assessed based on mesocosms and field experiments, rather than on population density (e.g. egg masses) and habitat alone. Furthermore, our study illustrates that planning through a collaborative effort between a developer and academic institution can reveal relevant site-specific scientific assessments to guide environmentally sensitive land development practices in a way that respects and possibly benefits the commercial interests of the developers. We hope, as in our case, these findings will ultimately inform planning and regulations to more adequately reflect current ecological understanding.

## Acknowledgement

I wish to thank Steward Pickett, Peter Morin, Steven Handel, Kristina Hill, Marsha Morin, Jeremy Feinberg, Chris Camacho, Tim Delorm, Brook Dannemiller, Brian Goldberg, Brian Field, Mike Sutton, Matt Dials, Renee Kaufman, Lia Kelerchian Tim Terway, Andrew Dance and the Related Company, Laura Newgard of EcolSci ences, Miele Construction, Donna Coogan and Paul Roggeman. Thanks to Janine Felson, Jacob Felson, Nancy Felson, Madeleine, Lev and Caroline.

